# NyxBind: enhancing DNN representations via contrastive learning for TFBS prediction

**DOI:** 10.1101/2025.10.21.683808

**Authors:** Xu Yang, Qingfa Xiao, Yucheng Xu, Jixin Yang, Yusen Hou, Weicai Long, Miaojun Huang, Yanlin Zhang

## Abstract

While pretrained genomic language models effectively capture general DNA sequence patterns through masked language modeling, they often struggle to discriminate subtle yet biologically critical differences among transcription factor binding site (TFBS) motifs. Recent studies suggest that contrastive learning can enhance the discriminative power of embeddings by explicitly modeling inter-instance similarities and differences. Building on this insight, we introduce NyxBind, the first TFBS prediction model that applies contrastive learning across multiple TFBS types to enhance regulatory sequence representations. NyxBind better captures discriminative sequence features, enabling more accurate and biologically meaningful TFBS prediction. Extensive evaluations show that NyxBind consistently outperforms alternative models across multiple TFBS classification benchmarks, demonstrating strong robustness and generalizability. Moreover, NyxBind supports both full-parameter and parameter-efficient fine-tuning while maintaining high performance, and supports accurate motif visualization, aligning closely with experimentally validated transcription factor binding profiles. The code are available at https://github.com/ai4nucleome/NyxBind.

## Introduction

Transcription factors (TFs) regulate gene expression by binding to short DNA regions known as transcription factor binding sites (TFBSs), typically 6–20 base pairs in length [1, 2]. Accurate identification of TFBSs is crucial for understanding gene regulation, interpreting non-coding variants, and studying cellular processes such as differentiation and disease progression [3, 4]. Each TF recognizes specific sequence motifs, commonly represented as position weight matrices (PWMs) and visualized with sequence logos. These motifs can be inferred from high-throughput assays such as ChIP-seq, SELEX, or PBMs [5], and identified computationally using tools like MEME and HOMER [6, 7]. However, TF binding specificity is influenced not only by core motifs but also by flanking sequence context and higher-order genomic features such as chromatin accessibility and DNA shape [8], necessitating more expressive models.

Deep learning has significantly advanced TFBS prediction by enabling the automatic learning of complex sequence features. DeepBind [9] pioneered the use of convolutional neural networks (CNNs) for this task, learning motif-like patterns directly from one hot encoded DNA sequences. This approach was extended by models such as DeepSEA [10], which employed deeper CNN architectures to predict a range of epigenomic features, and DanQ [11], which combined CNNs and recurrent neural networks to model both short and longrange dependencies.

More recently, inspired by advances in natural language processing, genomic language models (gLMs) have emerged as powerful tools for DNA sequence representation. DNABERT [12] adapts the BERT architecture [13] to DNA by tokenizing sequences into k-mers and pretraining on reference genomes using masked language modeling (MLM) objective. DNABERT2 [14] builds on this with architectural improvements and more efficient training strategies. Several other gLMs including Nucleotide Transformer [15], MutBERT [16], and GPN-MSA [17] further expand on this idea using diverse pretraining corpora and modeling choices. Additionally, models like Caduceus [18] and Evo [19] employ alternative architectures for efficient long-sequence modeling.

Despite their success in capturing general sequence patterns, gLMs pretrained solely with MLM are not optimized for tasks that require distinguishing between closely related sequence classes—such as identifying which transcription factor a given binding site belongs to. Although MLM enables rich contextual representation learning, it lacks an explicit objective to group functionally similar sequences and separate dissimilar ones in the embedding space. As a result, embeddings from MLM pretraining may be suboptimal for classification tasks that demand high discriminative power. To address this, several recent studies have explored augmenting MLM with additional training strategies. For instance, DNABERT-S [20] introduces contrastive learning to enhance cross-species generalization by enforcing separation between functionally distinct sequences; however, it yields little improvement in TFBS identification tasks. Other task-specific adaptation, such as BERT-TFBS [21] and BertSNR [22], combine MLM-pretrained models with convolutional layers, attention mechanisms, or multi-task learning frameworks to enhance TFBS prediction. While these models leverage the strengths of MLM, their performance gains highlight the need for explicit supervision at the representation level to better capture fine-grained regulatory distinctions.

Here, we introduce NyxBind, a contrastive learning framework that leverages multiple TFBS sequences to further pretrain existing MLM-based genomic models. NyxBind employs an advanced contrastive learning strategy to construct positive and negative sequence pairs across different TFBSs, capturing sequence-level similarities and differences and restructuring the embedding space to enhance functional separability. This approach enables the model to more effectively distinguish closely related TFBSs, producing embeddings that are both highly discriminative and biologically meaningful, and better suited for downstream regulatory tasks. By leveraging TFBS-derived supervision, NyxBind specifically addresses this limitation, achieving substantial gains in TFBS-specific tasks.

The main contributions of our study are as follows:

1. We propose, for the first time, a contrastive learning approach for TFBS classification.
2. We conduct comprehensive ablation studies across multiple training objectives.
3. We develop a motif visualization method that outperforms several existing deep learning approaches.
4. The proposed model achieves superior prediction performance compared with existing models, demonstrating strong results under both LoRA and full-parameter fine-tuning settings.

## Materials and methods

### Data

Contrastive learning was employed to further enhance the pretrained model using 690 human ChIP-seq TFBS datasets from the ENCODE project [23], all mapped to the hg19 reference genome, which were also utilized in previous models such as DeepSEA. These datasets comprise 160 TFs across 91 human cell types, with detailed infomation provided in Supplementary Table S1. To facilitate contrastive learning and because TFBS motifs of the same TF are generally conserved across cell types, sequences from the same TF but different cell lines were grouped into a single class, resulting in 160 labeled TFBS categories. For contrastive training, 101 bp sequences centered at ChIP-seq peak summits were extracted from all chromosomes except 11 and 12. Sequences from the same TF type were treated as positive pairs, while those from different TF types were treated as negative pairs. To ensure rigorous data partitioning and prevent information leakage, chromosomes 11 and 12 were initially merged into a single dataset, which was then randomly and evenly divided into validation and test subsets. The dataset construction and detailed information are described in Section 2.2.2. During contrastive learning, the model was trained to maximize representation similarity between sequences bound by the same TF (positive pairs) and minimize similarity between sequences bound by different TFs (negative pairs), thereby reinforcing the model’s ability to capture TF-specific binding patterns.

For TFBS prediction benchmarking (Section 2.3.3), 159 datasets were selected from the 690 ChIP-seq datasets, each containing more than 200 sequences on chromosomes 11 and 12 to ensure sufficient coverage for validation and testing. These datasets cover 33 human cell types and 35 TFs and represent commonly used datasets in models such as BERT-TFBS. Positive samples consisted of 101 bp sequences centered at ChIP-seq peak summits, while negative samples were drawn from non-overlapping regions matched for GC content, following the approach used in DNABERT2. A complete list of the datasets is provided in Supplementary Table S2.

For motif discovery (section 3.3), 33 TFBS datasets from the JASPAR database [24] were used. These datasets, previously employed in the BertSNR framework, cover a diverse range of transcription factor families, with each dataset corresponding to a specific TFBS. Positive sequences contained a single TFBS randomly positioned within the sequence, and negative samples were generated via dinucleotide-preserving shuffling using uShuffle [25]. This construction strategy was adopted to facilitate direct comparison with BertSNR. To enable independent motif visualization, an additional set of 33 JASPAR TFBS datasets not used during model training was employed. Detailed dataset composition is reported in Supplementary Table S3.

### NyxBind Overview

NyxBind is pre-trained through a two-stage process, as illustrated in Figure 1. In the first stage, MLM is used to pretrain a BERT-based model on large-scale genomic sequences. We adopt DNABERT2 as our backbone model, leveraging its robust and generalizable representations learned from multi-species genomic data. In the second stage, we apply contrastive learning to TFBS-associated sequences from different transcription factors. By constructing positive and negative sequence pairs based on shared or distinct TF motifs, this stage encourages the model to learn more discriminative representations that capture subtle regulatory differences.

**Fig. 1.**
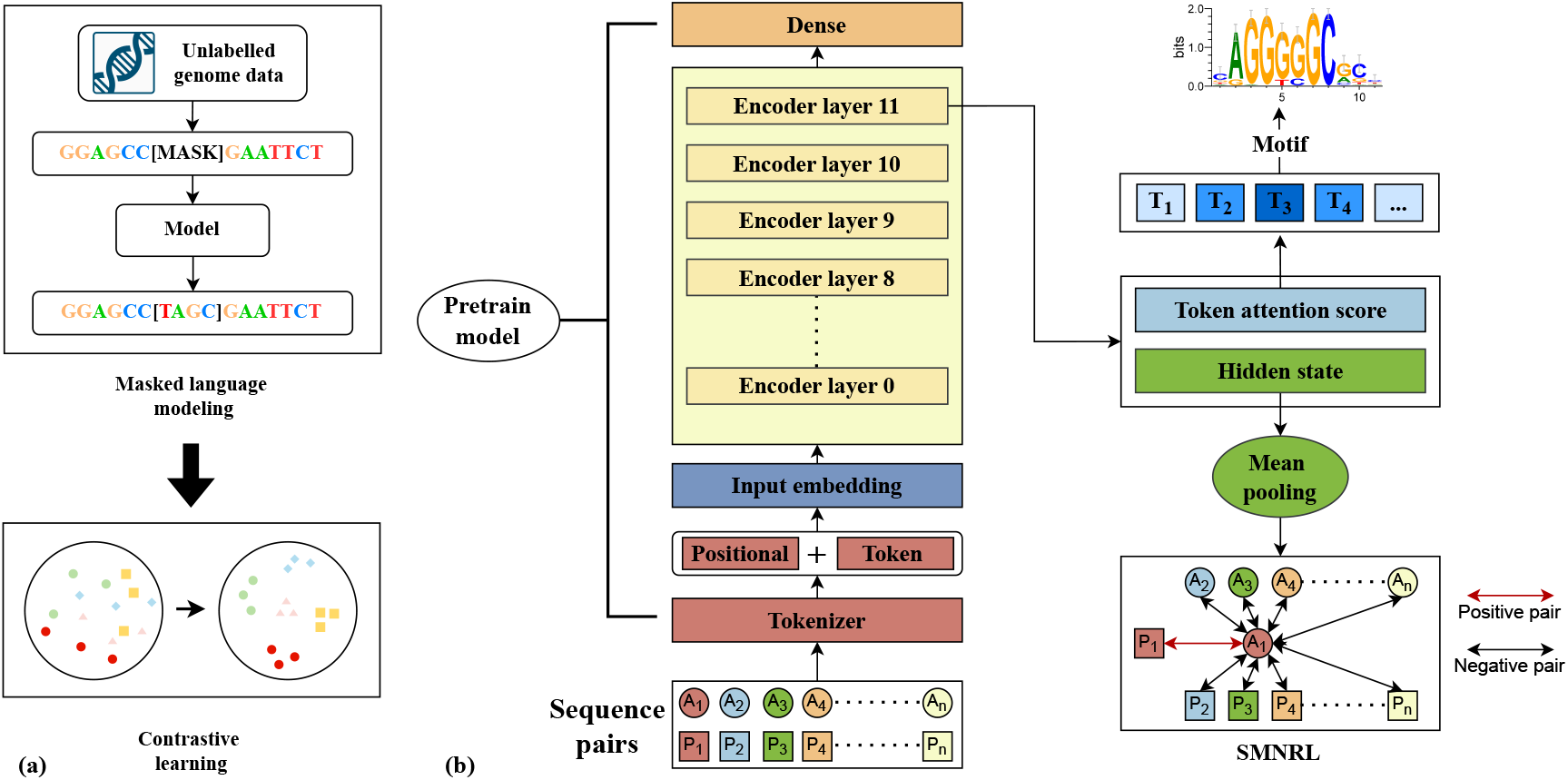
The framework of NyxBind. (a) NyxBind leverages the DNABERT2 model pretrained via MLM on multi-species genomes, and incorporates contrastive learning to improve discrimination across diverse TFBS types. (b) Brief demonstration of contrastive learning. The model is trained using the Symmetric Multiple Negatives Ranking Loss (SMNRL), where both directions of similarity — anchor-to-positive and positive-to-anchor — are jointly optimized to enhance representation consistency. Each batch contains N anchor–positive (A–P) pairs, where each pair belongs to the same TFBS type and forms a positive sample. For each anchor, positives from other pairs serve as in-batch negatives, and vice versa. Hidden states are extracted from the final encoder layer, followed by mean pooling to obtain sequence embeddings. In addition, NyxBind enables extraction of attention scores from each transformer layer, which can be used for motif visualization.

Following contrastive pretraining, NyxBind can be further fine-tuned on task-specific TFBS prediction datasets. We support both full-parameter fine-tuning and parameter-efficient adaptation using Low-Rank Adaptation (LoRA) [26], offering flexibility for deployment under varying computational constraints.

#### Model Structure

NyxBind adopts an architecture based on DNABERT2, combining a byte-pair encoding (BPE) tokenizer [27] with an embedding layer that integrates token and positional information. The BPE tokenizer segments DNA sequences into variable-length tokens by iteratively merging frequently co-occurring nucleotide pairs. This approach improves motif representation, shortens input length, reduces redundancy, and enhances computational efficiency.

The encoder consists of 12 Transformer layers with multi-head self-attention and several architectural optimizations. These include Flash Attention [28], an attention implementation that reduces memory access overhead and improves training speed; low-precision layer normalization for better computational efficiency; and the GEGLU activation function, which offers greater expressiveness compared to ReLU. A pooling layer is used to aggregate token-level outputs into a sequence-level embedding.

For downstream TFBS classification tasks, a task-specific classification head is added during fine-tuning. Both full-parameter and parameter-efficient fine-tuning modes are supported, allowing flexible adaptation of the model to different settings and datasets.

#### Contrastive learning

To enhance the discriminative capacity of the pretrained MLM-based model, NyxBind introduces an additional contrastive learning stage, employing Symmetric Multiple Negatives Ranking Loss (SMNRL) as the primary optimization objective. This supervised contrastive learning framework is inspired by SimCSE, originally proposed for sentence representation learning in NLP, and is adapted to the genomic domain using TFBS datasets.

To validate the effectiveness of SMNRL and assess alternative formulations, we conducted ablation experiments comparing four loss functions — SMNRL, MultipleNegativesRankingLoss (MNRL)[29], GISTEmbedLoss (GEL)[30] and ContrastiveLoss (CL)[31]. Among them, SMNRL consistently demonstrated the best overall discriminative performance, justifying its adoption as the final loss function in NyxBind.

During contrastive training, we utilized sequences containing TFBS collected from 690 ChIP-seq datasets, covering 160 distinct TFBS types. For the SMNRL, MNRL, and GEL objectives, 50,000 positive sequence pairs were constructed per TFBS type, resulting in a total of 8 million positive pairs. These objectives were trained without explicitly providing negative pairs, as negative pairs were implicitly formed within each batch. In contrast, for the CL objective, 25,000 positive pairs and 25,000 negative pairs were generated per TFBS type, also yielding 8 million pairs in total. Positive pairs were sampled from sequences of the same TFBS type, while negative pairs were constructed from sequences of different TFBS types to enhance inter-class discrimination. To prevent data leakage, sequences from chromosomes 11 and 12 were excluded from the contrastive pretraining stage. For validation and testing, we further constructed 500 positive and 500 negative sequence pairs per TFBS from sequences combined from chromosomes 11 and 12, resulting in 160,000 pairs for the validation set and 160,000 pairs for the test set. These sets were specifically designed to evaluate whether the model can produce distinguishable embeddings for different TFBSs.

To assess the utility of the contrastive pretraining for downstream tasks, we performed fine-tuning on 159 ChIP-seq datasets to conduct ablation experiments, enabling a fair comparison of different training objectives and their impact on task performance. All contrastive learning experiments were conducted for 3 epochs using a batch size of 128 and a learning rate of 3e-5. Training was performed on a single NVIDIA A800 GPU, and the complete process took approximately 12 hours.

#### Training Objective

We adopt SMNRL as the primary training objective for contrastive learning. SMNRL is a symmetric extension of MNRL, effective in settings where only positive pairs are available, such as semantically or functionally similar sequences. By computing the loss in both directions (anchor→positive and positive→anchor), SMNRL encourages robust and discriminative embeddings. This symmetric computation allows all remaining samples in the mini-batch—including anchors and positives from other pairs—to serve as implicit negatives.

Formally, given a batch of *N* positive sequence pairs (*a*_*i*_, *p*_*i*_), where *a*_*i*_ is the anchor and *p*_*i*_ the corresponding positive, let *s*(*·, ·*) denote a similarity function (e.g., cosine similarity), and *τ* a temperature hyperparameters. The SMNRL objective is:

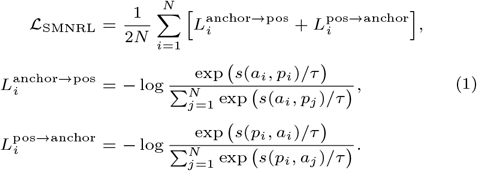

To evaluate the effectiveness of SMNRL, we perform ablation studies using alternative objectives:

- **MultipleNegativesRankingLoss** computes the loss solely in the anchor→positive direction, where all other positives in the mini-batch are treated as implicit negatives, while anchors from other pairs are excluded:

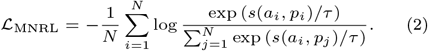
- **GISTEmbedLoss** dynamically selects informative negatives using a guide model. For each anchor-positive pair 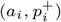, candidates more similar than the reference positive *p*_*i*_ are treated as positives 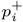, and less similar ones as negatives 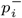. Here, the similarity scores are denoted as 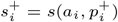 for the positive pair, and 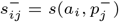 for the guide-selected negatives:

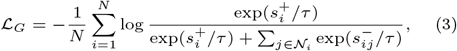

where 𝒩_*i*_ is the set of guide-selected negatives.
- **ContrastiveLoss** computes the cosine similarity between embeddings of a sentence pair (*a, p*) produced by a `SentenceTransformer` model, and compares it to the target label *y ∈* {0, 1} using the classic contrastive loss. Here, the hyperparameter margin (default 0.5) specifies the minimum distance for negative pairs. The loss is defined as:

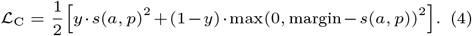

### Evaluation

#### Evaluation metrics

For contrastive learning, we evaluate the quality of the learned representations using the binary classification evaluator module from the Sentence Transformers framework. This evaluator computes cosine similarity between sequence pairs and reports three metrics: cosine average precision (cosine AP), cosine F1 score, and cosine accuracy. Cosine AP measures the model’s ability to rank positive pairs above negative ones across a range of thresholds, providing a threshold-independent evaluation. Cosine F1 and cosine accuracy are computed using a fixed similarity threshold, capturing performance trade-offs between precision, recall, and overall classification accuracy.

TFBS prediction is typically formulated as a binary classification task [32]. To evaluate model performance, we consider four widely used classification metrics: accuracy (ACC), ROC-AUC, PR-AUC, and Matthews Correlation Coefficient (MCC). ACC is a threshold-based metric that measures the proportion of correctly classified samples and provides a basic assessment of overall performance. ROC-AUC, a threshold-free metric, evaluates the model’s ability to discriminate between positive and negative classes across varying decision thresholds. PR-AUC, also threshold-free, focuses on the trade-off between precision and recall and is particularly informative in imbalanced datasets where the positive class is sparse. MCC is a threshold-based metric that combines all four confusion matrix components (true/false positives and negatives) into a single score, offering a balanced evaluation even in the presence of class imbalance. Together, these metrics capture both overall and class-specific performance under different evaluation perspectives, providing a comprehensive assessment of TFBS prediction accuracy.

#### Fine-Tuning Strategies for NyxBind

NyxBind supports two strategies for adapting the pretrained model to TFBS prediction tasks: full-parameter fine-tuning and parameter-efficient fine-tuning via Low-Rank Adaptation (LoRA).

Full-parameter fine-tuning updates all model weights, offering maximal task adaptation but is computationally intensive. As an efficient alternative, LoRA fine-tuning freezes the pretrained weights and updates only low-rank adapter parameters, reducing trainable parameters while retaining task-specific learning capacity

To compare the effectiveness of these two fine-tuning strategies, we performed both full-parameter and LoRA based fine-tuning across 159 TFBS classification tasks. For full-parameter fine-tuning, we apply a learning rate of 3e-5, batch size of 32, 5 training epochs. For LoRA, we apply a learning rate of 7e-4, batch size of 32, 5 training epochs, LoRA rank of 8, LoRA alpha of 16, and a dropout rate of 0.05.

#### Evaluation of NyxBind and alternative Models for TFBS Prediction

We selected a diverse set of representative models spanning both traditional architectures and pretrained transformer-based approaches. DeepBind employs a single-layer CNN with one-hot encoding to capture local sequence motifs. DanQ builds on this architecture by combining CNNs with bidirectional long short-term memory (Bi-LSTM) layers to model both local and long-range dependencies. We used the PyTorch implementations provided in DeepSTF [32], with a learning rate of 1e-3, batch size of 64, and 15 training epochs, following the hyperparameters settings reported in the original paper. The NT series [15] includes several large-scale transformer models pretrained on different genomic datasets. NT-500M-Human was trained on the human reference genome (hg38), NT-2500M-1000G on 3,202 human genomes from the 1000 Genomes Project, and NT-2500M-Multi on 850 genomes from multiple species. NT-v2-500M-Multi incorporates architectural refinements including rotary positional embeddings, SwiGLU activations, and the removal of biases and dropout layers. All NT models were fine-tuned using LoRA with a learning rate of 1e-4, batch size of 32, 5 training epochs, LoRA rank of 8, LoRA alpha of 16, and a dropout rate of 0.05, following the hyperparameters settings used in DNABERT2 [14]. DNABERT2 is a BERT-based genomic language model pretrained on multi-species genomic sequences, leveraging ALiBi positional encoding and FlashAttention to improve scalability and training efficiency. Conversely, DNABERTS focuses on producing species-aware DNA embeddings, enabling better cross-species representation learning. BERT-TFBS extends DNABERT2 by integrating CNN layers and a Convolutional Block Attention Module (CBAM), enabling enhanced motif detection. For DNABERT2, DNABERTS, and NyxBind, we followed the hyperparameters settings of DNABERT2, used a learning rate of 3e-5, batch size of 32, except that the number of training epochs was increased to 5. For BERT-TFBS, we followed the hyperparameters settings reported in the original paper, using a learning rate of 1.5e-5, batch size of 32, and 15 training epochs to ensure convergence given its added complexity. Transformer-based models often serve as embedding backbones in hybrid architectures. To evaluate the generalization ability of NyxBind in this context, we replaced the DNABERT2 encoder in BERT-TFBS with NyxBind and fine-tuneing using same hyperparameters, yielding a variant referred to as BERT-TFBS_N. This experiment assesses whether the representations learned by NyxBind retain their effectiveness when transferred to a downstream architecture.

All models were optimized using the AdamW optimizer with a weight decay of 0.01 and 30 warm-up steps.These settings were adopted across all models, as preliminary experiments showed that variations in these hyperparameterss had minimal impact on performance.In addition, early stopping based on validation loss was applied to prevent overfitting and to retain the model at near-optimal validation performance.

### Motif Analysis

To assess whether NyxBind captures biologically meaningful sequence patterns, we developed a motif visualization framework adapted from DNABERT [12]. The goal is to extract transcription factor binding motifs learned by the model and evaluate their alignment with experimentally validated reference motifs.

We obtained attention weights from the final Transformer layer of the NyxBind model, specifically from the [CLS] token to all other tokens. Each token’s attention score was redistributed across its corresponding nucleotide positions, producing a base-resolution attention map. These scores were standardized using z-score normalization to generate a quantitative attention profile along the input sequence. A sliding window was then applied to this profile to identify high-confidence regions. Windows with aggregate scores exceeding a predefined threshold were selected, and the central regions were extracted as candidate motif instances. These instances were aligned to produce position weight matrices (PWMs) and sequence logos, enabling direct comparison with canonical TF motifs.

To evaluate motif recovery performance, we followed the experimental setup proposed in BertSNR [22]. We first fine-tuned each model on 33 human TFBS datasets from the JASPAR database. To avoid data leakage during evaluation, we then applied motif discovery to a separate set of 33 matched datasets corresponding to the same transcription factors. This pairing ensured that the evaluation was conducted on held-out sequences not seen during training, while maintaining biological relevance through 1-to-1 TF correspondence.

We compared NyxBind against three baselines for motif visualization: (1) BertSNR, a BERT-based model that supports single-nucleotide-resolution prediction; (2) DeepSNR [33], a hybrid CNN–deconvolutional model; (3) D-AEDNet [34], an encoder–decoder architecture that integrates convolutional and attention layers to enhance feature extraction and information flow. Motif similarity was assessed using the TOMTOM tool [35], which compares discovered motifs against all human TFBS PWMs in the JASPAR database. Statistical significance of motif matches was evaluated using p-values, e-values, and q-values. The p-value represents the probability of observing a match score by chance, the e-value estimates the expected number of false positives in the database search, and the q-value is the false discovery rate–adjusted p-value accounting for multiple testing. Matches were considered significant if they met all of the following thresholds: p-value < 0.1, e-value < 0.5, and q-value < 0.01. Model performance was measured by the number of significantly matched motifs across 33 evaluation datasets.

## Results

### Training Objective Variants

We perform the ablation study on SMNRL and other contrasitive learning methods include MNRL, GEL and CL. For SMNRL, MNRL and GEL, training will only include positive pairs. For a batch N, with a pair of anchor sequence and positive sequence, SMNRL will consider N-1 other anchors and N-1 postives as negative samples, MNRL will consider N-1 positives as negative samples. While GEL use a guide model, which we use DNABERT2, to eval 2N-2 sequences by calculate the similarity of embedding generated by guide model to decide whether to be negative samples or positive samples for the anchor.

We evaluated these four methods on 159 ChIP-seq datasets for the classification task. As shown in Table 1, SMNRL achieves the highest MCC, which attributed to the use of more in-batch pairs compared to MNRL and CL. In contrast, the relatively lower performance of GEL could be due to the quality of embeddings generated by the guide model.

**Table 1.** Ablation study on training objective variants. Values indicate the MCC (%) difference relative to SMNRL on 159 ChIP-seq datasets.

| Training Objective | MCC Difference (%) |
| --- | --- |
| SMNRL | 77.27 |
| MNRL | -0.10 |
| CL | -0.36 |
| GEL | -1.66 |

### Evaluation

#### Contrastive Learning Enhances NyxBind Representations for TFBS

We employed SMNRL to train our model with the aim of improving its capacity to capture meaningful and discriminative representations of sequences across diverse TFBS ChIP-seq datasets. This strategy encourages the model to map similar sequences closer in the embedding space, while pushing dissimilar sequences apart. The final model obtained after training is referred to as NyxBind.

To assess NyxBind’s ability to generate informative representations of TFBS, we evaluated its performance in producing discriminative embeddings for pairs of similar and dissimilar TFBS sequences on test sets, which is shown at Table 2. The evaluation was based on cosine similarity between embedding vectors, using metrics includes cosine AP, cosine accuracy, and cosine F1 score. NyxBind exhibits substantial improvements over the baseline, with a 26.01% increase in cosine AP, a 15.75% increase in cosine accuracy, and a 3.50% increase in cosine F1. These results highlight the effectiveness of contrastive learning in enhancing the model’s ability to generate consistent embeddings for functionally similar sequences and to distinguish between different TFBS binding patterns.

**Table 2.** Performance comparison between DNABERT2 and NyxBind using cosine-based evaluation metrics.

| Model | Cosine AP | Cosine Accuracy | Cosine F1 |
| --- | --- | --- | --- |
| DNABERT2 | 0.5327 | 0.5428 | 0.6667 |
| NyxBind | **0.7928** | **0.7003** | **0.7017** |

#### Attention Visualization Reveals Motif Focus

To further investigate the impact of contrastive learning across diverse TFBS datasets, we visualized the attention maps of NyxBind before and after contrastive pretraining. We randomly selected sequences associated with CTCFL, FOXA1, FOSL1, and GATA2 with motifs centered in the sequences for attention visualization. In these examples, NyxBind shows attention concentration on motif-associated tokens in some layers compared with the baseline model. Here, the attention weights were computed as the average across all heads in each layer, providing a layer-wise view of motif-focused attention. The complete visualization results for these TFs are provided in Supplementary Figure S1.

As illustrated in Figure 2, we randomly selected a sequence from the FOSL1 dataset (MA0477.2.5, JASPAR) and visualized attention distributions across all layers. In the early layers, attention is primarily concentrated on the [CLS] and [SEP] tokens, reflecting the model’s capture of global sequence structure and boundaries, a pattern also observed in BertSNR. After contrastive learning, the attention in the deeper layers gradually shifts toward tokens corresponding to the FOSL1 motif. In Figure 2a (before contrastive learning), the attention remains relatively diffuse with no clear focus on motif regions, whereas Figure 2b (after contrastive learning) shows dense and highly focused attention lines on motif-associated tokens in the later layers, highlighting that NyxBind places greater emphasis on motif regions when generating sequence embeddings.

**Fig. 2.**
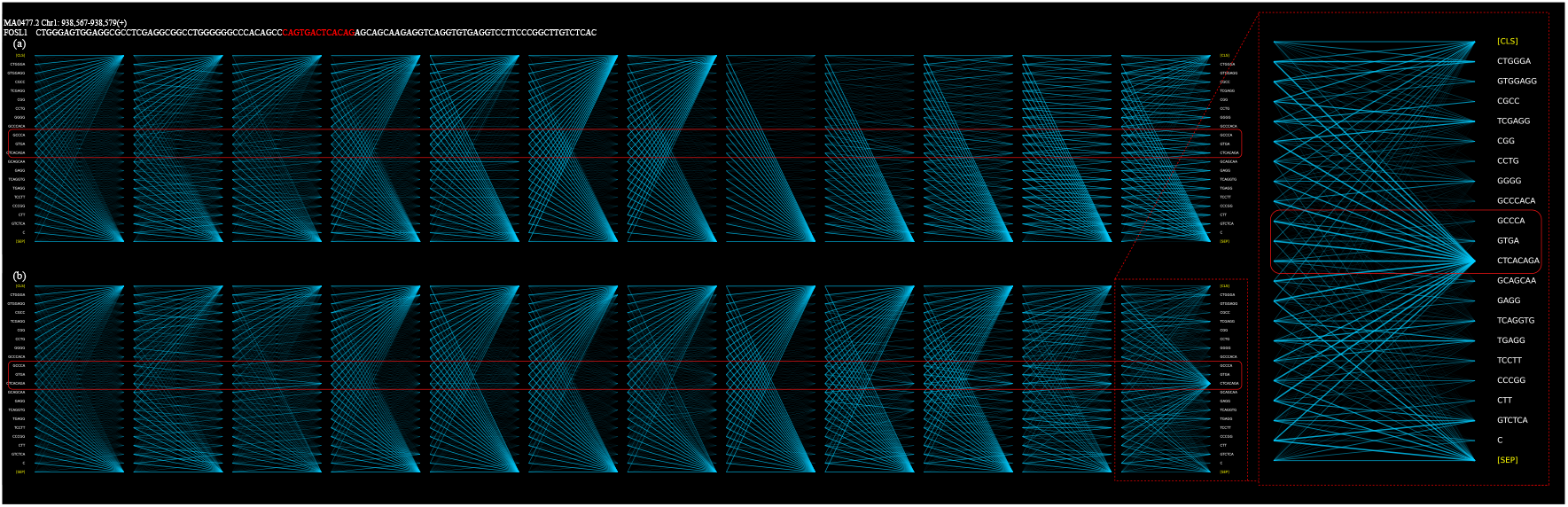
Attention visualization before and after contrastive learning. Visualization of attention weights across all layers for a sequence sampled from the FOSL1 TFBS dataset (MA0477.2, JASPAR). (a) Model before contrastive learning (baseline) shows relatively diffuse attention across the sequence. (b) Model after contrastive pretraining (NyxBind) exhibits strong and focused attention centered on the FOSL1 motif, indicating improved motif recognition.

#### NyxBind improves TFBS prediction

We conducted a comprehensive comparison between our model and nine other methods—DeepBind, DanQ, DNABERT2, DNABERTS, BERT-TFBS, NTv2-500M-Multi, NT-2500M-Multi, NT-500M-Human, and NT-2500M-1000G—across 159 ChIP-seq datasets for TFBS prediction. Detailed results are provided in Supplementary Table S4. As shown in Table 3 and Figure 3(a), our model consistently outperforms alternative methods across all evaluation metrics—including MCC, ACC, PR-AUC, and ROC-AUC, averaged over 159 TFBS ChIP-seq datasets. NyxBind outperforms the second-best model, DNABERT2, achieving absolute gains of 2.33% in ACC, 4.71% in MCC, 1.25% in PR-AUC, and 1.51% in ROC-AUC. As illustrated in the scatter plot of Figure 3(b), NyxBind exhibits robust and consistent improvements across all four evaluation metrics on most of the 159 ChIP-seq datasets. These results highlight the superior performance and generalization ability of our approach for TFBS prediction tasks. This demonstrates the strong capability of NyxBind in capturing regulatory sequence features and highlights its potential as a robust foundation for a wide range of downstream applications.

**Table 3.** Benchmark results of models on 159 TFBS ChIP-seq datasets.

| Model | ACC | MCC | PR-AUC | ROC-AUC |
| --- | --- | --- | --- | --- |
| **NyxBind** | **0.8888** | **0.7727** | **0.9557** | **0.9485** |
| DNABERT2 | 0.8655 | 0.7256 | 0.9432 | 0.9334 |
| DNABERTS | 0.8652 | 0.7248 | 0.9428 | 0.9329 |
| NTv2-500M-Multi | 0.8623 | 0.7189 | 0.9400 | 0.9289 |
| NT-2500M-Multi | 0.8582 | 0.7111 | 0.9386 | 0.9271 |
| NT-500M-Human | 0.8092 | 0.6104 | 0.9005 | 0.8858 |
| NT-2500M-1000G | 0.8379 | 0.6697 | 0.9210 | 0.9100 |
| DeepBind | 0.8340 | 0.6601 | 0.9185 | 0.9022 |
| DanQ | 0.8305 | 0.6538 | 0.9172 | 0.8993 |
| BERT-TFBS | 0.8565 | 0.7076 | 0.9371 | 0.9260 |
| BERT-TFBS_N | 0.8848 | 0.7656 | 0.9543 | 0.9459 |

**Fig. 3.**
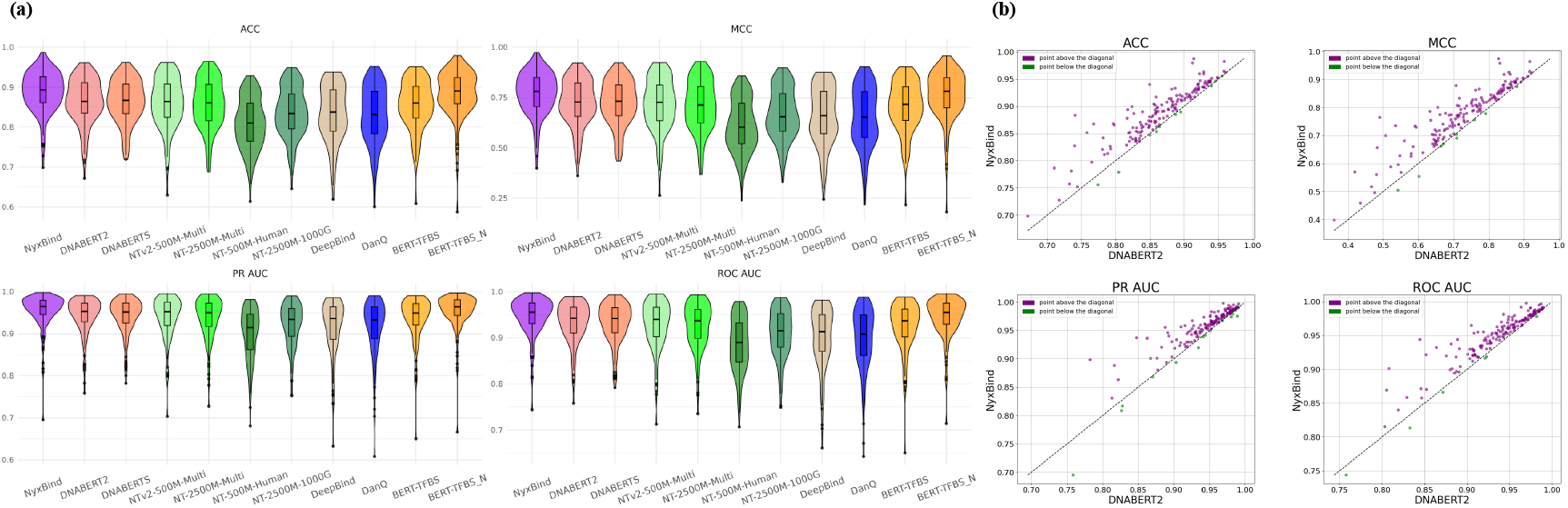
Evaluation results on 159 ChIP-seq datasets. (a) Violin plots compare NyxBind with baseline models across 159 datasets, showing the distribution of MCC, ACC, ROC-AUC, and PR-AUC scores. Each violin plot contains an embedded box plot indicating the median and interquartile range. (b) Scatter plots comparing ACC, MCC, PR-AUC, and ROC-AUC metrics between NyxBind and DNABERT2 across the 159 datasets, with each point representing a dataset.

We evaluated DNABERTS on TFBS prediction and found its performance largely comparable to DNABERT2 across most datasets. This aligns with DNABERTS’ limitation: it performs contrastive learning across multi-species sequences rather than TFBS-specific sequences, making it less specialized for TFBS discrimination.

To further evaluate the effectiveness of NyxBind as a sequence embedding model, we conducted additional experiments by replacing the DNABERT2 encoder in BERT-TFBS with NyxBind, while keeping all other components unchanged—including the CNN, CBAM, and the output layers. The resulting configuration, referred to as BERT-TFBS_N, was fine-tuned using full-parameter training on 159 ChIP-seq datasets. As shown in Table 3, BERT-TFBS_N consistently outperforms the original BERT-TFBS model across all evaluation metrics. These results demonstrate that NyxBind provides more informative embeddings than a model only pretrained via MLM and enhances downstream TFBS prediction when integrated into hybrid architectures.

#### NyxBind performs well with both LoRA and full-parameter fine-tuning

We fine-tune both NyxBind and DNABERT2 on 159 TFBS ChIP-seq datasets using two strategies: full-parameter fine-tuning and parameter-efficient fine-tuning via LoRA. As shown in Table 4, NyxBind maintains consistently high performance across both tuning strategies, with only marginal differences between FT and LoRA: 0.21% in ACC, 0.47% in MCC, 0.05% in PR-AUC, and 0.13% in ROC-AUC. These results demonstrate NyxBind’s strong adaptability to diverse fine-tuning paradigms. In contrast, DNABERT2 exhibits more performance drop when switching from FT to LoRA, with declines of 1.69% in ACC, 3.46% in MCC, 1.55% in PR-AUC, and 1.69% in ROC-AUC, indicating lower robustness to parameter-efficient adaptation. As shown in Figure 4, each point in the scatter plots represents a single TFBS ChIP-seq dataset, comparing performance between full-parameter fine-tuning (FT) and LoRA across four metrics: ACC, MCC, PR-AUC, and ROC-AUC. These comparisons reveal that NyxBind consistently achieves more stable and higher performance than DNABERT2 under both fine-tuning strategies.

**Table 4.**
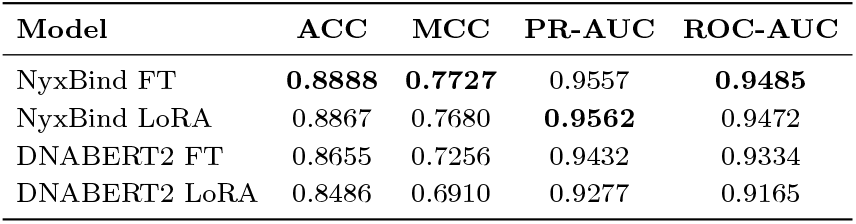
Comparison of LoRA and FT performance on 159 ChIP-seq datasets.

**Fig. 4.**
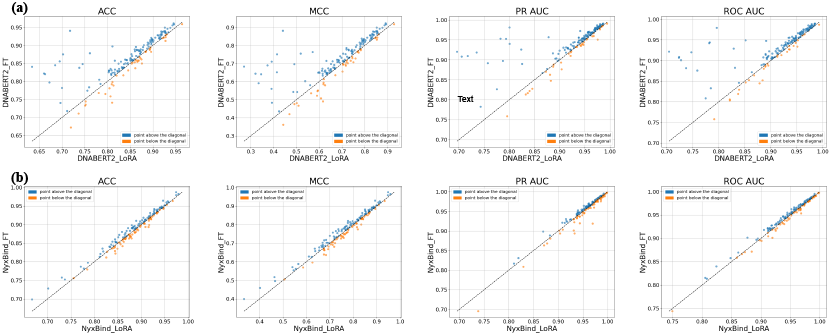
Scatter plots comparing FT (y-axis) and LoRA (x-axis) across ACC, MCC, PR-AUC, and ROC-AUC metrics. The diagonal line denotes equal performance on 159 ChIP-seq datasets. (a) Results of DNABERT2. (b) Results of NyxBind.

### NyxBind Effectively Identifies Motifs from Sequences

In this study, we further examine NyxBind’s ability to highlight biologically relevant sequence motifs using the model’s self-attention mechanism. We extracted attention scores at the token level from the final Transformer layer. Since each token spans multiple nucleotides because of BPE, we computed nucleotide-level attention by dividing the token-level score equally among the nucleotides it represents. The resulting base-resolution scores were then standardized using z-score normalization and used for visualization of attention-derived motifs.

To ensure fair evaluation and avoid data leakage, NyxBind was fine-tuned on 33 TFBS datasets from the JASPAR database, and motif discovery was performed on a matched, held-out set of 33 TFBS datasets representing the same transcription factors. For visualization, the attention-derived motifs were extracted using a fixed window size of 11. We generated PWMs from the attention scores and compared them to all human TFBS PWMs in the JASPAR database using TOMTOM. NyxBind recovered the largest number of matching motifs, identifying 31 out of 33 TFBS patterns. BertSNR followed with 30 matches, while both DeepSNR and D-AEDNet correctly identified 17 motifs each. Although NyxBind recovered the largest number of motifs, BertSNR produced slightly more statistically significant alignments according to p-value distributions, which can be attributed to BertSNR’s multi-task fine-tuning strategy, in contrast to NyxBind’s fine-tuning method. Detailed motif-level comparisons are provided in Supplementary Table S5 and Table S6.

Figure 5 shows example motif logos for four TFs (FOXA1, CTCFL, FOSL1, and USF1), generated by NyxBind and three baseline models, alongside the corresponding JASPAR reference motifs. NyxBind accurately reconstructs core sequence features and demonstrates strong motif fidelity, reinforcing its capacity not only for accurate TFBS prediction but also for revealing biologically meaningful sequence patterns associated with transcription factor binding

**Fig. 5.**
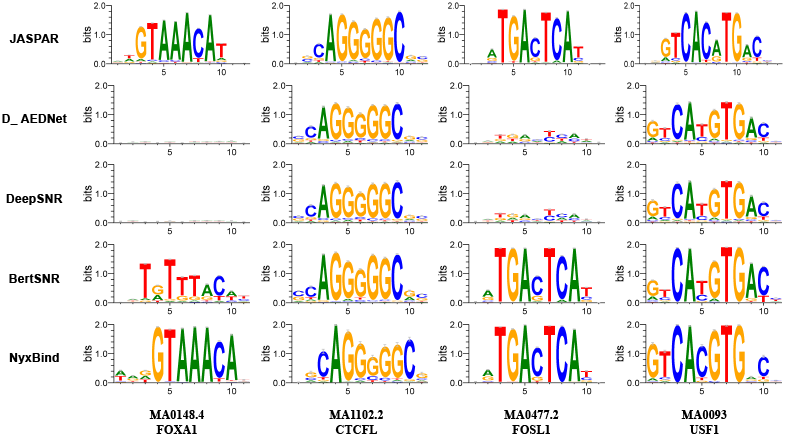
Motif visualization for four TFBS types—FOXA1, CTCFL, FOSL1, and USF1. For each TFBS, we present the reference motif from JASPAR alongside motifs predicted by D_AEDNet, DeepSNR, BertSNR, and NyxBind. A predicted motif is considered successful if it closely resembles the reference or captures complementary binding patterns.

## Discussion

In this study, we presented NyxBind, a contrastive learning framework designed to refine sequence representations for TFBS prediction. Through large-scale benchmarking on 159 ChIP-seq datasets, NyxBind consistently outperformed existing models across diverse evaluation settings, demonstrating strong generalization across transcription factors, cell types, and binding contexts While existing genomic foundation models are typically pretrained using self-supervised objectives such as MLM or next-token prediction, these objectives are not explicitly designed to discriminate between functionally distinct regulatory sequences. NyxBind addresses this limitation by applying contrastive learning to TFBS-labeled sequences, explicitly pulling together sequences from the same TF and pushing apart those from different TFs. This task-aligned supervision encourages the model to learn more discriminative and functionally meaningful representations, leading to substantial performance improvements in TFBS classification.

Although NyxBind shows strong performance in TFBS prediction and motif visualization, future work could focus on expanding the training datasets with additional transcription factors, integrating cross-species regulatory sequences, and introducing more detailed similarity annotations to further improve the model’s robustness and biological generalization.

In summary, we demonstrate a contrastive learning approach that enhances the representation of TFBSs. By explicitly modeling similarities and differences between regulatory sequences, NyxBind generates highly predictive representations, providing a solid foundation for further developments in regulatory genomics.

### Key Points

- NyxBindis an advanced model that enhances DNA sequence representations for TFBS prediction through contrastive learning across 160 TF types.
- The proposed SMNRL strategy significantly improves embedding quality, leading to higher cosine similarity–based metrics and better TFBS prediction performance.
- NyxBind exhibits novel capabilities in motif visualization, accurately highlighting biologically meaningful binding regions.
- Evaluations on 159 ChIP-seq datasets show that NyxBind consistently outperforms other deep learning models in TFBS prediction.

## SUPPLEMENTARY DATA

Supplementary data are available online at http://bib.oxford/journals.org/

## Acknowledgments

This work extends a course project from DSAA 6000M: *ML in Genetics & Genomics* at HKUST(GZ). The scientific computations were performed on the High Performance Computing Platform at HKUST (GZ) with support from the College of Future Technology.

