## Supplementary Figure∼S1 for "NyxBind: enhancing DNN representations via contrastive learning for TFBS prediction"

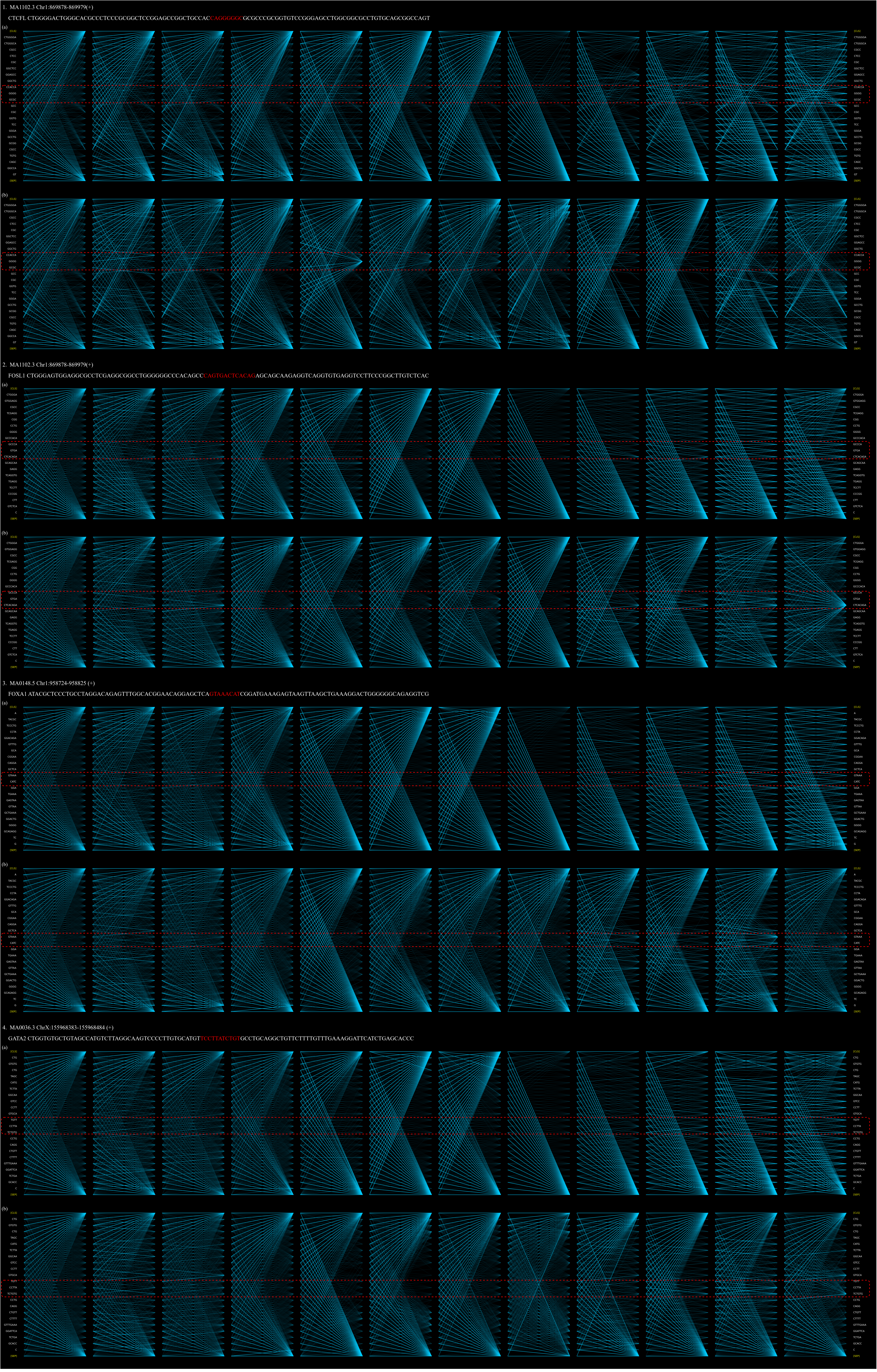


**Supplementary Figure S1. Attention visualization before and after contrastive learning.**

We randomly selected sequences associated with CTCFL, FOXA1, FOSL1, and GATA2, each containing motifs centered within the sequence, for attention visualization. These line plots illustrate the attention weight distribution across all layers for each token during inference. Compared with the pre-contrastive learning model (a), the post-contrastive learning model (b) assigns noticeably higher attention weights to motif-related tokens in some layers, indicating improved motif awareness after contrastive learning.
